# Selection and co-selection of antibiotic resistances among *Escherichia coli* by antibiotic use in primary care: an ecological analysis

**DOI:** 10.1101/573360

**Authors:** Koen B Pouwels, Berit Muller-Pebody, Timo Smieszek, Susan Hopkins, Julie V Robotham

**Author notes:** Correspondence to: Koen B. Pouwels, Modelling and Economics Unit, National Infection Service, Public Health England, 61 Colindale Ave, London NW9 5EQ, United Kingdom.

## Abstract

The majority of studies that link antibiotic usage and resistance focus on simple associations between the resistance against a specific antibiotic and the use of that specific antibiotic. However, the relationship between antibiotic use and resistance is more complex. Here we evaluate which antibiotics, including those mainly prescribed for respiratory tract infections, are associated with increased resistance among *Escherichia coli* isolated from urinary samples.

Monthly primary care prescribing data were obtained from National Health Service (NHS) Digital. Positive *E. coli* records from urine samples in English primary care (n=888,207) between April 2014 and January 2016 were obtained from the Second Generation Surveillance System. Elastic net regularization was used to evaluate associations between prescribing of different antibiotic groups and resistance against amoxicillin, cephalexin, ciprofloxacin, co-amoxiclav and nitrofurantoin at the clinical commissioning group (CCG) level. England is divided into 209 CCGs, with each NHS practice prolonging to one CCG.

Amoxicillin prescribing (measured in DDD/ 1000 inhabitants / day) was positively associated with amoxicillin (RR 1.03, 95% CI 1.01 – 1.04) and ciprofloxacin (RR 1.09, 95% CI 1.04 – 1.17) resistance. In contrast, nitrofurantoin prescribing was associated with lower levels of resistance to amoxicillin (RR 0.92, 95% CI 0.84 – 0.97). CCGs with higher levels of trimethoprim prescribing also had higher levels of ciprofloxacin resistance (RR 1.34, 95% CI 1.10 – 1.59).

Amoxicillin, which is mainly (and often unnecessarily) prescribed for respiratory tract infections is associated with increased resistance against various antibiotics among *E. coli* causing urinary tract infections. Our findings suggest that when predicting the potential impact of interventions on antibiotic resistances it is important to account for use of other antibiotics, including those typically used for other indications.

**Author summary:** Antibiotic resistance is increasingly recognised as a threat to modern healthcare. Effective antibiotics are crucial for treatment of serious bacterial infections and are necessary to avoid that complicated surgical procedures and chemotherapy becoming life-threatening. Antibiotic use is one of the main drivers of antibiotic resistance. The majority of antibiotic prescriptions are prescribed in primary care, however, a large proportion of these antibiotic prescriptions are unnecessary. Understanding which antibiotics are causing antibiotic resistance to what extent is needed to prevent under- or over-investment in interventions lowering use of specific antibiotics, such as rapid diagnostic tests for respiratory tract infection.

We have statistically evaluated which antibiotics are associated with higher and lower levels of antibiotic resistance against common antibiotics among *Escherichia coli* bacteria sampled from the urinary tract by comparing antibiotic prescribing and resistance in different geographical areas in England. Our model shows that amoxicillin, the most commonly used antibiotic in England and mainly used for respiratory tract infections, is associated with increased resistance against several other antibiotics among bacteria causing urinary tract infections. The methods used in this study, that overcome several of the limitations of previous studies, can be used to explore the complex relationships between antibiotic use and antibiotic resistance in other settings.

## Introduction

In England, approximately three-quarters of antibiotics are dispensed in primary care [1]. A substantial proportion of these antibiotics are unnecessary, being used for viral or self-limiting respiratory tract infections [2, 3]. When antibiotics are used for a viral infection an effect on the pathogen causing the infection, both in terms of outcome of the infection as well as resistance against antibiotics, is not expected. However, because antibiotics typically used for respiratory tract infections, such as amoxicillin, have a systemic effect, they can select for antibiotic resistances among bacteria that are carried by the host at the moment of treatment, i.e. bacteria forming the microflora or microbiota [4]. If those bacteria are pathogenic or act as a reservoir of resistance elements this may lead to an increased incidence of symptomatic infections caused by bacteria that are resistant to clinically important antibiotics [5, 6]. Moreover, antibiotic prescriptions are often longer than necessary, which could further increase antibiotic resistance levels without clinical benefit [7]. However, the relationship between antibiotic use and antibiotic resistance is more complex. There may be cross-resistance between antibiotics, such as observed for ampicillin and amoxicillin [8]. Resistance genes may be linked on the same mobile genetic element, such as observed for amoxicillin and trimethoprim resistance genes [8, 9]. Therefore treatment with one antibiotic may select for resistance against another antibiotic via cross-resistance and co-selection [8, 9]. Treatment with one antibiotic may also simply kill competing bacterial flora, thereby providing bacteria resistant to another antibiotic more space and nutrients, such as anti-anaerobic antibiotics that promote the overgrowth of vancomycin-resistant enterococci [10, 11]. Moreover, mutations or acquired genes conferring resistance to one antibiotic can not only increase but also decrease resistance to another antibiotic [12]. Such collateral sensitivity, where resistance against one antibiotic confers sensitivity against another has been mainly explored for spontaneous resistance mutations [12, 13].

The vast majority of studies that link antibiotic usage and resistance at the population level focus on simple associations between the resistance against a specific antibiotic and the use of that specific antibiotic or antibiotic group, or alternatively group all antibiotics together [14]. There is a lack of studies that simultaneously take into account use of different antibiotics and potential co-selection.

We therefore evaluated associations between prescribing levels of antibiotic groups in primary care in England and resistance against amoxicillin, cephalexin, ciprofloxacin, co-amoxiclav and nitrofurantoin, among *Escherichia coli* isolated from urinary samples in England, thereby taking into account prescribing of other antibiotics groups. Because we only had data on antibiotic prescribing in primary care, we focused on *E. coli* sampled from the urinary tract by general practitioners. We used elastic net regularization [15, 16], because this method – which combines the advantages of both least absolute shrinkage and selection operator (lasso) [17] and ridge regression [18] – works particularly well in situations with high collinearity and relative large number of variables compared to the amount of observations [15, 16]. This is particularly relevant, because there are many different antibiotic groups and there are likely strong correlations between prescribing patterns of antibiotics leading to sparsity and multicollinearity problems with standard regression techniques [19].

The vast majority urinary tract infections are caused by *E. coli* infections and uropathogenic *E. coli* strains are often part of the human intestinal microflora. Given the systemic nature of systemic antibiotics, this research may shed light on the question whether and to what extent antibiotics typically being used to treat (viral) respiratory tract infections, such as amoxicillin [1], may result in resistance problems against not only the same antibiotic, but also other antibiotics among bacteria for which the antibiotic courses were not initially intended.

The work presented in this paper provides evidence about which antibiotics are associated with higher and lower levels of antibiotic resistance against common antibiotics among *Escherichia coli* bacteria sampled from the urinary tract by comparing antibiotic prescribing and resistance in different geographical areas in England. Our models show that amoxicillin, the most commonly used antibiotic in England and mainly used for respiratory tract infections, is associated with increased resistance against several other antibiotics among bacteria causing urinary tract infections. The methods used in this study, that overcome several of the limitations of previous studies, can be used to explore the complex relationships between antibiotic use and antibiotic resistance in other settings.

## Results

The antibiotic groups that were used most intensively with ≥1 daily defined doses (DDD) per 1000 inhabitants per day, were tetracyclines, penicillins with extended spectrum (mainly amoxicillin) [1], macrolides, Beta-lactamase-resistant penicillins (mainly <u>Flucloxacillin</u>) [1], and trimethoprim (Fig 1).

**Fig 1.**
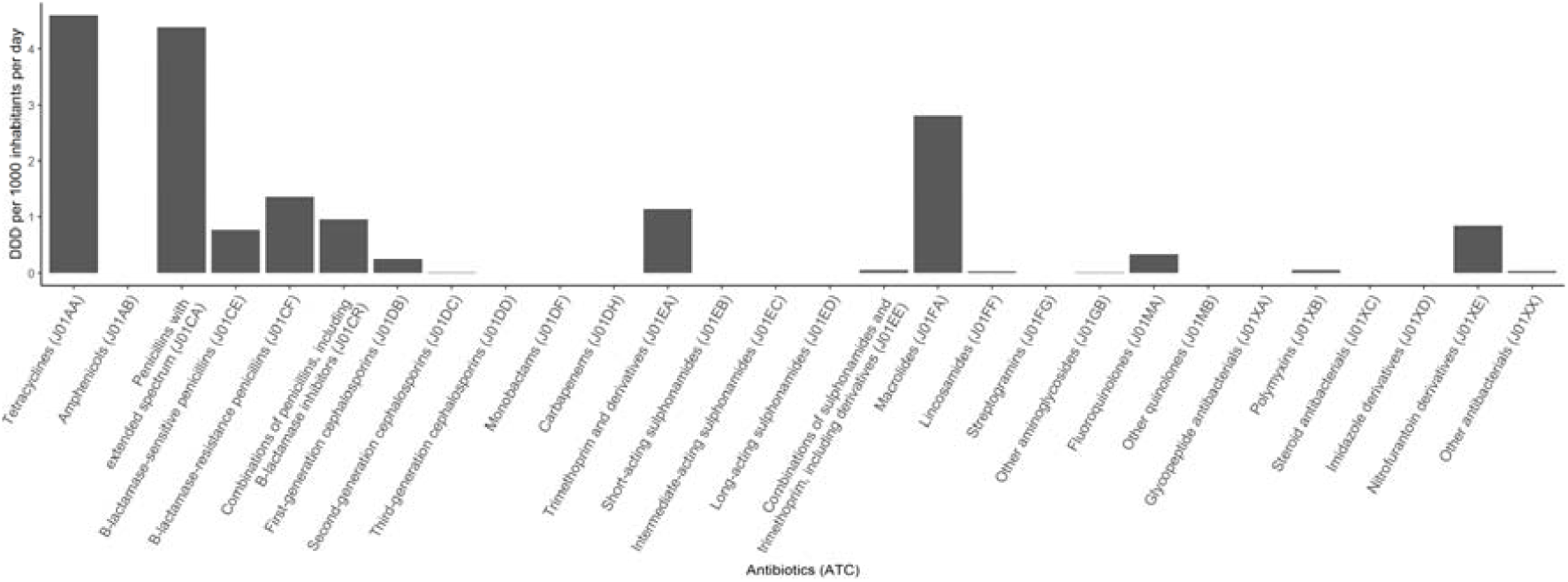
The average DDD per 1000 inhabitants per day for different antibiotic groups during the study period.

While we evaluated the association between resistances of interest and all antibiotic groups, Fig 2 shows the variation in prescribing between the different clinical commissioning groups (CCGs) for the 4 antibiotic groups that are prescribed the most. In addition, these maps show the variation in nitrofurantoin and trimethoprim, which are the antibiotics typically used to treat urinary tract infections. There was substantial variation in antibiotic prescribing between the different CCGs (Fig 2), some CCGs had high antibiotic prescribing levels for all antibiotics, especially in the North of England. There was generally more variation in antibiotic prescribing between CCGs than variation over time within CCGs. However, for some antibiotics there were clear peaks in the amount of dispensed antibiotics, in line with peaks in the incidence of respiratory tract infections (winter) and skin infections (summer) (S1, Fig S1-S2).

**Fig 2.**
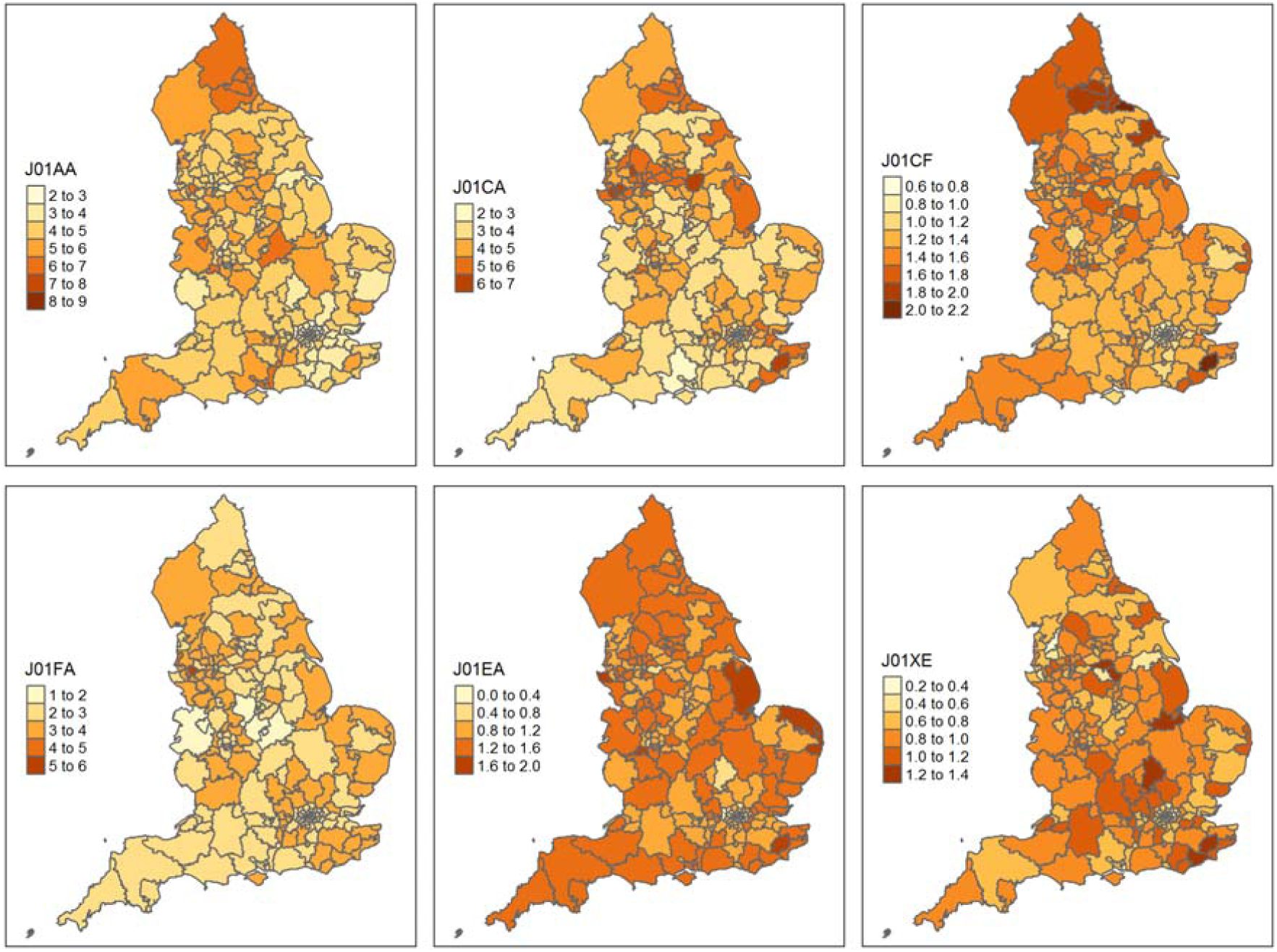
Maps of the average number of DDD per 1000 inhabitants per day for the 209 clinical commissioning groups during the study period. Not that different scales are used for the different antibiotics. J01AA = tetracyclines; J01CA = penicillins with extended spectrum (mainly amoxicillin); J01CF = Beta-lactamase-resistant penicillins (mainly flucloxacillin); J01FA = macrolides; J01EA = trimethoprim; J01XE = nitrofurantoin.

Between April 2014 and January 2016, nearly all (99%, n=888,207) *E. coli* urinary samples from general practice patients sent in for laboratory testing were tested for resistance against nitrofurantoin. The percentages of samples tested for resistance against the other included antibiotics varied between 78% for amoxicillin and 90% for co-amoxiclav.

There was substantial variation in the percentage of *E. coli* urinary isolates that were resistant to the antibiotics tested (Table 1).

**Table 1.**
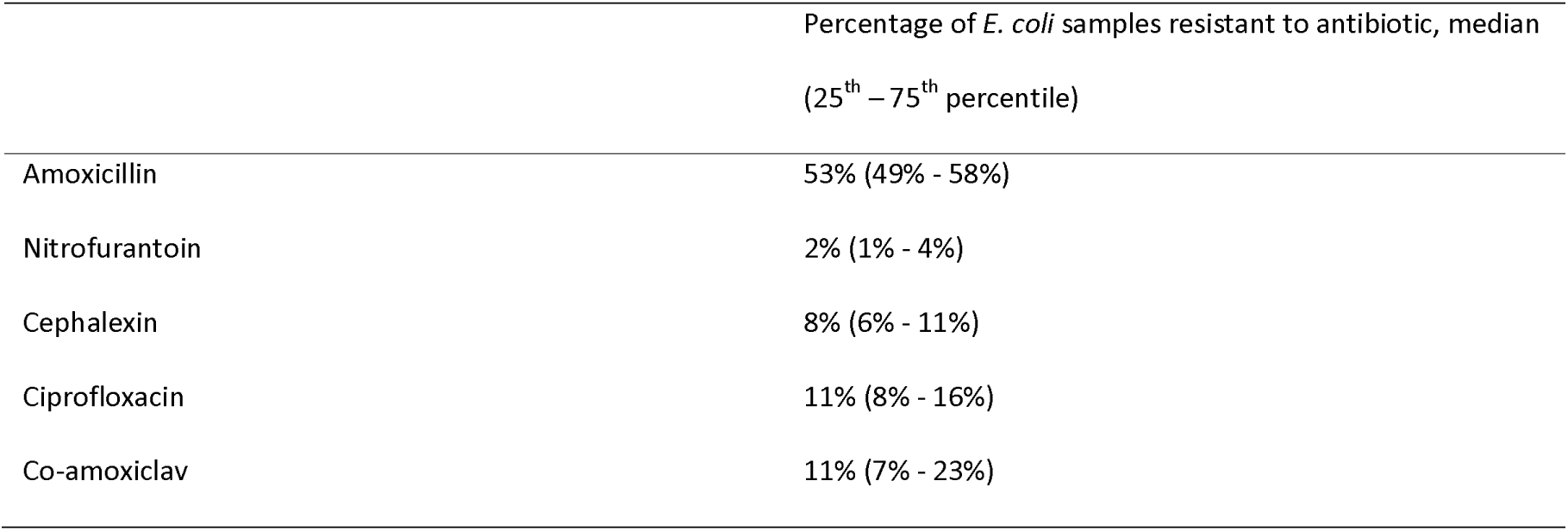
Variation in antibiotic resistance among *E. coli* urinary samples, measured on a monthly basis at the clinical commissioning group level.

|  | Percentage of *E. coli* samples resistant to antibiotic, median<br>(25<sup>th</sup> – 75<sup>th</sup> percentile) |
| --- | --- |
| Amoxicillin | 53% (49% - 58%) |
| Nitrofurantoin | 2% (1% - 4%) |
| Cephalexin | 8% (6% - 11%) |
| Ciprofloxacin | 11% (8% - 16%) |
| Co-amoxiclav | 11% (7% - 23%) |

There was less variation in the percentage of isolates that were resistant to the antibiotics test over time (S1, Fig S3-S7). The variation in testing rate, which may influence apparent antibiotic resistance proportions, and variation in the measured antibiotic resistance proportions are shown in Fig 3. As is apparent from the maps, part of the variation in the apparent proportion of samples that are resistant to antibiotics can be explained by the test rate. When few tests are determined, most of the samples are resistant. However, there are also regions with a relatively high test rate and still relatively high resistance, such as in the North-East, indicating that the resistance prevalence may indeed be relatively high.

**Fig 3.**
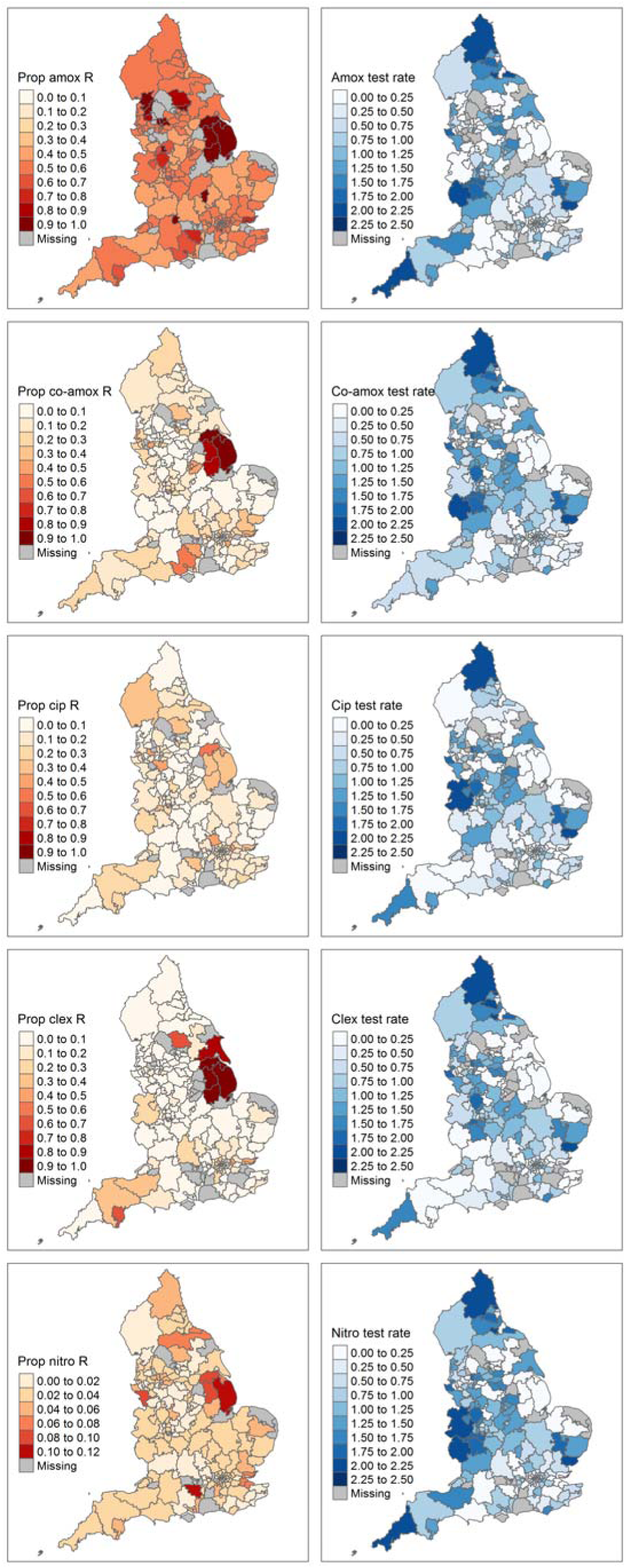
The left column shows the proportion of *E. coli* urinary samples that are resistant to amoxicillin, co-amoxiclav, ciprofloxacin, cephalexin and nitrofurantoin, respectively. The right column shows the number of samples tested for resistance against these antibiotics per 1000 person-months.

Results from the elastic net regularization models showed amoxicillin resistance was positively associated with prescribing of penicillins with extended spectrum (mainly amoxicillin in England)^1^ in the month (RR 1.03, 95%CI 1.01 to 1.04), quarter (RR 1.03, 95%CI 1.01 to 1.04) and year (RR 1.04, 95%CI 1.01 to 1.06) (Table 2; the full results including the coefficients for the test rate are provided in S2) before the specimen date.

**Table 2.** Associations between amoxicillin resistance among *E. coli* urinary samples and antibiotic prescribing (DDD per 1000 persons per day)

| Antibiotic prescribed | Amoxicillin resistance, antibiotic prescribing 1 month before. RR (2.5th–97.5th percentile of bootstrap) | Amoxicillin resistance, antibiotic prescribing 1-3 month before. RR (2.5th–97.5th percentile of bootstrap) | Amoxicillin resistance, antibiotic prescribing 1 year before. RR (2.5th–97.5th percentile of bootstrap) |
| --- | --- | --- | --- |
| Tetracyclines (J01AA) | 1.00 (0.98 – 1.01) | 1.00 (0.98 – 1.02) | 1.00 (0.98 – 1.04) |
| Penicillins with extended spectrum (J01CA) | **1.03 (1.01 – 1.04)<sup>a</sup>** | **1.03 (1.01 – 1.04)<sup>a</sup>** | **1.04 (1.01 – 1.06)<sup>a</sup>** |
| Beta-lactamase-sensitive penicillins (J01CE) | 1.02 (0.97 – 1.12) | 1.00 (1.00 – 1.19) | - |
| Beta-lactamase-resistant penicillins (J01CF) | 1.03 (0.98 – 1.12) | 1.03 (0.96 – 1.13) | 1.04 (0.95 – 1.17) |
| Combinations of penicillins, including β-lactamase inhibitors (J01CR) | 1.02 (0.95 – 1.08) | - | - |
| First-generation cephalosporins (J01DB) | 1.01 (0.91 – 1.07) | - | - |
| Second-generation cephalosporins (J01DC) | 1.00 (0.85 – 1.02) | - | - |
| Trimethoprim and derivatives (J01EA) | 1.01 (0.98 – 1.08) | 1.01 (1.00 – 1.12) | 1.00 (1.00 – 1.17) |
| Macrolides (J01FA) | 0.99 (0.97 – 1.02) | 1.00 (0.97 – 1.03) | 1.00 (0.97 – 1.04) |
| Lincosamides (J01FF) | 0.98 (0.71 – 1.00) | - | - |
| Fluoroquinolones (J01MA) | **0.93 (0.78 – 0.99)<sup>a</sup>** | 0.87 (0.69 – 1.00) | 0.88 (0.61 – 1.00) |
| Polymyxins (J01XB) | 1.01 (0.93 – 1.35) | - | - |
| Nitrofuran derivatives (J01XE) | **0.92 (0.84 – 0.97)<sup>a</sup>** | **0.91 (0.82 – 0.96)<sup>a</sup>** | **0.91 (0.83 – 0.98)<sup>a</sup>** |
| Other antibacterials (J01XX) | 0.98 (0.87 – 1.13) | - | - |

<sup>a</sup>Associations for which 2.5<sup>th</sup> and 97.5<sup>th</sup> percentile of the clustered bootstrap are both indicating an increased or decreased risk.

A similar direct association was seen in that CCGs that used more nitrofurantoin had a higher percentage of *E. coli* samples that tested resistant to nitrofurantoin (RR 1.52, 95%CI 1.00 to 2.24) (Table 3). The data did not confirm such a relationship between first-generation cephalosporin use (mainly cephalexin in England)[1] and cephalexin resistance, between fluoroquinolone use (mainly ciprofloxacin in England)[1] and ciprofloxacin resistance, or between combinations of penicillins, including β-lactamase inhibitors (mainly co-amoxiclav in England)[1] and co-amoxiclav resistance (Tables 4-6).

**Table 3.**
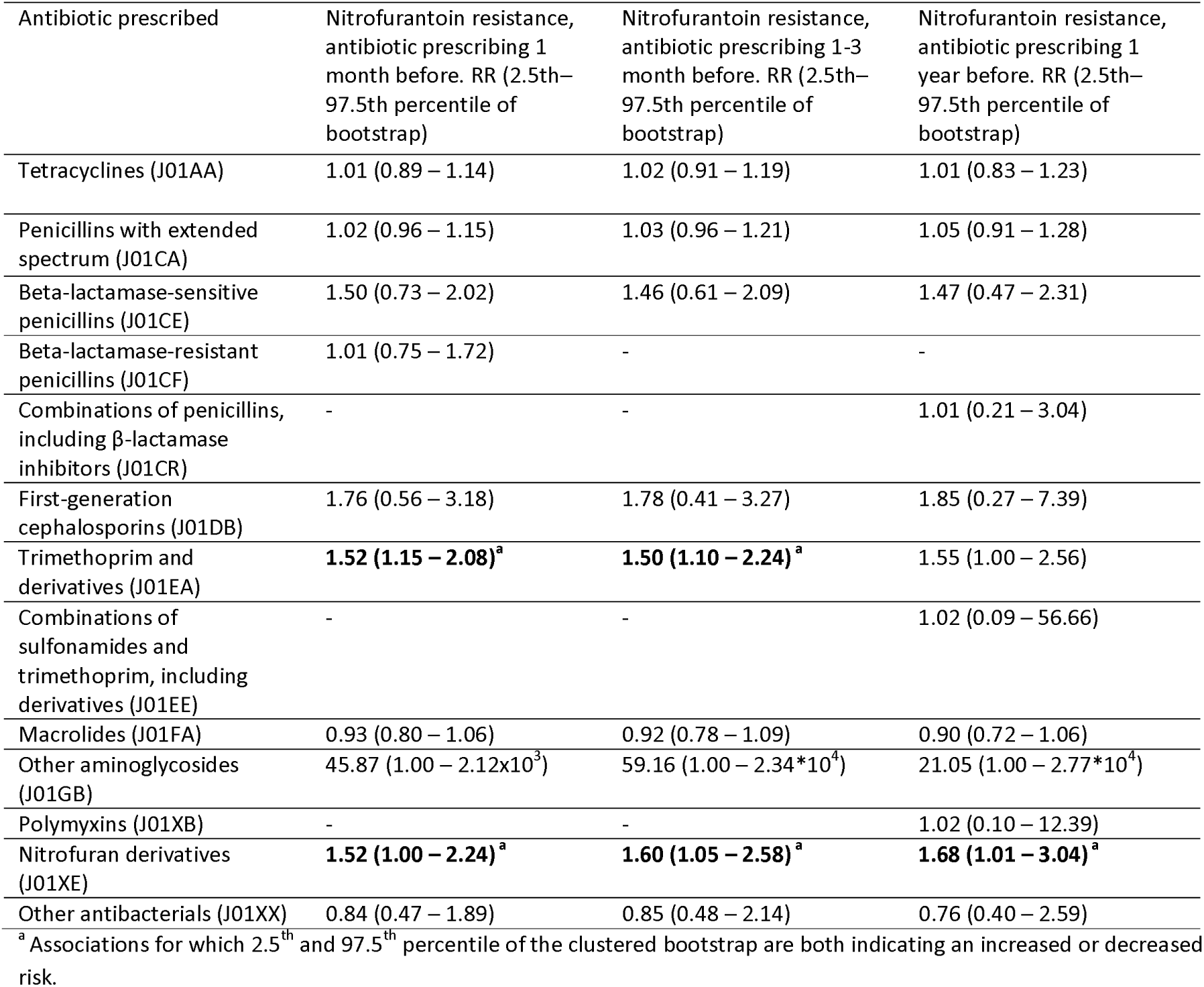
Associations between nitrofurantoin resistance among *E. coli* urinary samples and antibiotic prescribing (DDD per 1000 persons per day)

| Antibiotic prescribed | Nitrofurantoin resistance, antibiotic prescribing 1 month before. RR (2.5th–97.5th percentile of bootstrap) | Nitrofurantoin resistance, antibiotic prescribing 1-3 month before. RR (2.5th–97.5th percentile of bootstrap) | Nitrofurantoin resistance, antibiotic prescribing 1 year before. RR (2.5th–97.5th percentile of bootstrap) |
| --- | --- | --- | --- |
| Tetracyclines (J01AA) | 1.01 (0.89 – 1.14) | 1.02 (0.91 – 1.19) | 1.01 (0.83 – 1.23) |
| Penicillins with extended spectrum (J01CA) | 1.02 (0.96 – 1.15) | 1.03 (0.96 – 1.21) | 1.05 (0.91 – 1.28) |
| Beta-lactamase-sensitive penicillins (J01CE) | 1.50 (0.73 – 2.02) | 1.46 (0.61 – 2.09) | 1.47 (0.47 – 2.31) |
| Beta-lactamase-resistant penicillins (J01CF) | 1.01 (0.75 – 1.72) | - | - |
| Combinations of penicillins, including β-lactamase inhibitors (J01CR) | - | - | 1.01 (0.21 – 3.04) |
| First-generation cephalosporins (J01DB) | 1.76 (0.56 – 3.18) | 1.78 (0.41 – 3.27) | 1.85 (0.27 – 7.39) |
| Trimethoprim and derivatives (J01EA) | **1.52 (1.15 – 2.08)<sup>a</sup>** | **1.50 (1.10 – 2.24)<sup>a</sup>** | 1.55 (1.00 – 2.56) |
| Combinations of sulfonamides and trimethoprim, including derivatives (J01EE) | - | - | 1.02 (0.09 – 56.66) |
| Macrolides (J01FA) | 0.93 (0.80 – 1.06) | 0.92 (0.78 – 1.09) | 0.90 (0.72 – 1.06) |
| Other aminoglycosides (J01GB) | 45.87 (1.00 – 2.12x10<sup>3</sup>) | 59.16 (1.00 – 2.34*10<sup>4</sup>) | 21.05 (1.00 – 2.77*10<sup>4</sup>) |
| Polymyxins (J01XB) | - | - | 1.02 (0.10 – 12.39) |
| Nitrofurantoin derivatives (J01XE) | **1.52 (1.00 – 2.24)<sup>a</sup>** | **1.60 (1.05 – 2.58)<sup>a</sup>** | **1.68 (1.01 – 3.04)<sup>a</sup>** |
| Other antibacterials (J01XX) | 0.84 (0.47 – 1.89) | 0.85 (0.48 – 2.14) | 0.76 (0.40 – 2.59) |

<sup>a</sup> Associations for which 2.5<sup>th</sup> and 97.5<sup>th</sup> percentile of the clustered bootstrap are both indicating an increased or decreased risk.

**Table 4.**
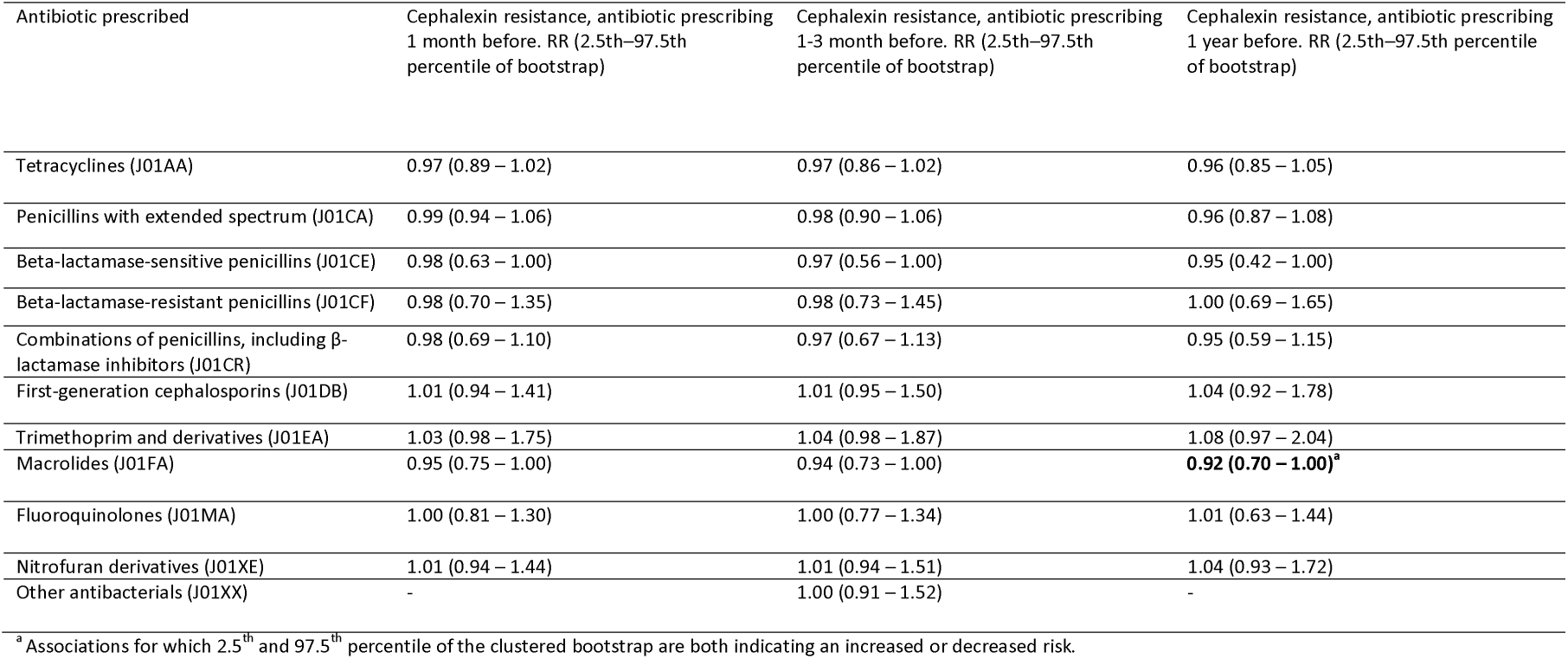
Associations between cephalexin resistance among *E. coli* urinary samples and antibiotic prescribing (DDD per 1000 persons per day)

**Table 5.** Associations between ciprofloxacin resistance among *E. coli* urinary samples and antibiotic prescribing (DDD per 1000 persons per day)

| Antibiotic prescribed | Ciprofloxacin resistance, antibiotic prescribing 1 month before. RR (2.5th–97.5th percentile of bootstrap) | Ciprofloxacin resistance, antibiotic prescribing 1-3 month before. RR (2.5th–97.5th percentile of bootstrap) | Ciprofloxacin resistance, antibiotic prescribing 1 year before. RR (2.5th–97.5th percentile of bootstrap) |
| --- | --- | --- | --- |
| Tetracyclines (J01AA) | **0.92 (0.88 – 0.98)<sup>a</sup>** | **0.92 (0.87 – 0.98)<sup>a</sup>** | 0.93 (0.85 – 1.01) |
| Penicillins with extended spectrum (J01CA) | **1.09 (1.04 – 1.17)<sup>a</sup>** | **1.10 (1.03 – 1.19)<sup>a</sup>** | 1.13 (1.02 – 1.25) |
| Beta-lactamase-resistant penicillins (J01CF) | - | - | 0.96 (0.65 – 1.24) |
| Trimethoprim and derivatives (J01EA) | **1.34 (1.10 – 1.59)<sup>a</sup>** | **1.35 (1.13 – 1.72)<sup>a</sup>** | **1.35 (1.11 – 1.78)<sup>a</sup>** |
| Macrolides (J01FA) | **0.85 (0.76 – 0.94)<sup>a</sup>** | **0.84 (0.75 – 0.94)<sup>a</sup>** | **0.84 (0.73 – 0.95)<sup>a</sup>** |
| Fluoroquinolones (J01MA) | 1.24 (1.00 – 2.81) | 1.29 (1.00 – 3.55) | 1.38 (1.00 – 4.40) |
| Nitrofurantoin | - | - | 1.00 (0.65 – 1.14) |

<sup>a</sup> Associations for which 2.5<sup>th</sup> and 97.5<sup>th</sup> percentile of the clustered bootstrap are both indicating an increased or decreased risk.

**Table 6.** Associations between co-amoxiclav resistance among *E. coli* urinary samples and antibiotic prescribing (DDD per 1000 persons per day)

| Antibiotic prescribed | Co-amoxiclav resistance, antibiotic prescribing 1 month before. RR (2.5th–97.5th percentile of bootstrap) | Co-amoxiclav resistance, antibiotic prescribing 1-3 month before. RR (2.5th–97.5th percentile of bootstrap) | Co-amoxiclav resistance, antibiotic prescribing 1 year before. RR (2.5th–97.5th percentile of bootstrap) |
| --- | --- | --- | --- |
| Tetracyclines (J01AA) | 1.03 (0.89 – 1.31) | 1.04 (0.87 – 1.33) | 1.04 (0.87 – 1.35) |
| Penicillins with extended spectrum (J01CA) | 0.99 (0.74 – 1.00) | 0.98 (0.70 – 1.00) | 0.97 (0.68 – 1.02) |
| Beta-lactamase-sensitive penicillins (J01CE) | - | - | 1.00 (1.00 – 8.86) |
| Combinations of penicillins, including β-lactamase inhibitors (J01CR) | - | - | 1.00 (0.51 – 2.62) |
| Trimethoprim and derivatives (J01EA) | - | - | 1.00 (0.54 – 2.30) |

However, it should be noted that a substantial proportion of these specific antibiotics are used in the hospital settings, for which no data was available [20]. Besides the obvious associations between prescribing of a particular antibiotic and resistance to that same antibiotic, we also observed associations between prescribing of a particular antibiotic and resistance against an antibiotic from another group. Amoxicillin use was not only associated with higher levels of amoxicillin resistance, but also with increased ciprofloxacin resistance (RR 1.09 95%CI 1.04 to 1.17) (Table 5) and increased trimethoprim resistance (as we have previously shown [8]). CCGs with high prescribing of trimethoprim also had higher levels of nitrofurantoin resistance (RR 1.52 95%CI 1.15 to 2.08) and ciprofloxacin resistance (RR 1.34 95%CI 1.10 to 1.59) (Tables 3 and 5).

There were also some antibiotics that had negative associations with antibiotic resistances. Nitrofurantoin use was associated with decreased amoxicillin resistance (RR 0.92 95%CI 0.84 to 0.97) (Table 2). Previously, we observed a similar negative association between nitrofurantoin use and trimethoprim resistance levels [8]. Tetracycline and macrolide use was associated with decreased ciprofloxacin resistance (Table 5), while fluoroquinolone use was associated with lower amoxicillin resistance levels (Table 2).

Results were very similar when restricting the analyses to months with at least 20 measurements (S2, Table S6-S10), suggesting that random error or remaining systematic error due to low testing probability were not large after adjusting for the test rate.

## Discussion

We found evidence of both selection and co-selection, as well as geographical patterns in antibiotic use and resistance. Amoxicillin use, an antibiotic that is mainly used for respiratory tract infections (∼83%) and rarely for urinary tract infections (∼2%) [1], is associated with increased resistance against amoxicillin and ciprofloxacin among urinary tract infections caused by *E. coli*. Areas that used more trimethoprim had higher levels of ciprofloxacin and nitrofurantoin resistance among *E. coli* urinary samples. These positive associations between prescribing of a particular antibiotic and resistance against another antibiotic suggest that co-selection may play a role.

We found that use of amoxicillin and trimethoprim were associated with resistance against ciprofloxacin, which suggests that co-selection may be occurring. Isolates from the common *E. coli* urinary pathogenic clonal group ST131 are often non-susceptible to both fluoroquinolones and trimethoprim-sulfamethoxazole and/or β-lactam antibiotics [21, 22], which may explain why use of trimethoprim and amoxicillin would select for ciprofloxacin resistance as amoxicillin use likely also selects for bacteria with trimethoprim resistance genes [8]. This link is further supported by the ECO·SENS study that found that resistance to any agent was correlated with increased resistance to all other agents tested, except for fosfomycin [9].

Nitrofurantoin had a negative association with amoxicillin resistance. This is in line with our previous finding that areas with relatively high nitrofurantoin use have lower trimethoprim resistance levels.^7^ Nitrofurantoin resistance genes are, in contrast to trimethoprim and amoxicillin resistance genes, not frequently found on mobile genetic elements with multiple resistances or correlated with multiple resistances in other ways [8, 11, 23]. Therefore, nitrofurantoin use may select for *E. coli* that are susceptible to amoxicillin and trimethoprim. Collateral sensitivity (and collateral resistance) has previously been observed for resistance against several antibiotics among *E. coli* isolates in an experimental setting [13]. Selecting for resistance against ampicillin was associated with increased sensitivity to nitrofurantoin compared to wild type *E. coli* strains [13], suggesting that collateral sensitivity may partly explain the observed negative association between amoxicillin and trimethoprim.

Given the high fitness cost of nitrofurantoin resistance [23], the positive association between trimethoprim use and nitrofurantoin resistance is not likely due to co-selection, but may be due to the possibility that CCGs with high trimethoprim usage have more patients on long-term treatment or prophylaxis with trimethoprim and nitrofurantoin.

The negative association between prescribing of tetracyclines or macrolides and ciprofloxacin resistance is harder to explain. Macrolides are typically active against Gram-positive bacteria, although they can be effective against Gram-negative bacteria when used in combination with antibiotics that do have outer-membrane disruptive activity [24]. Given the lack of selective pressure, *E coli* are unlikely to frequently harbor resistance mechanisms against antibiotics like macrolides, which could make such a synergistic combination therapy particularly effective against Gram-negative bacteria resistant to multiple antibiotics including ciprofloxacin [24]. However, this is unlikely the cause of the negative association between macrolide use and ciprofloxacin resistance, as such a combination therapy is not frequently being used or necessary in England. Collateral sensitivity has been observed for resistance against azithromycin (a macrolide) and sensitivity to nalidixic acid (a quinolone) among a pathogenic *E. coli* strain [13]. This may partly explain why macrolides use had a negative association with ciprofloxacin resistance. However, further studies are needed to evaluate whether these associations are causal.

Amoxicillin is the most frequently used antibiotic for respiratory conditions which are responsible for the largest share in inappropriate antibiotic prescribing in primary care [1-3]. Based on the current and a previous study, amoxicillin prescribing appears to be associated with increased resistance to amoxicillin, ciprofloxacin and trimethoprim [8]. Together these findings suggest that there is a substantial potential to reduce selective pressure via (co-)selection through reduction in the amount of unnecessary treatment with amoxicillin.

In many countries nitrofurantoin has been adopted as the first-line treatment for uncomplicated urinary tract infections [25,26]. Compared to other European countries, such as the Netherlands, the proportion of urinary tract infections treated with nitrofurantoin is much lower in England, though between June 2017 and June 2018 the ratio of trimethoprim prescribing over trimethoprim plus nitrofurantoin prescribing substantially decreased from 0.53 to 0.38, subsequent to the national quality premium [1, 26].

Recent recommendations are to prescribe nitrofurantoin as the first choice treatment for uncomplicated urinary tract infections [8]. We found nitrofurantoin prescribing to be associated with lower levels of resistance to amoxicillin and it has a negative association with trimethoprim resistance [8]. Conversely, we previously showed trimethoprim prescribing to be associated with increased resistance to trimethoprim [8], and, here, associated with high ciprofloxacin resistance. Our findings therefore suggest that a shift towards more nitrofurantoin instead of trimethoprim for uncomplicated urinary tract infections could potentially reduce antibiotic resistance among *E. coli*.

Patterns of co-resistance and co-selection likely differ between various parts of the world as there is substantial variation in selection pressure by antibiotics and infection prevention and control between countries [27,28]. However, we would even caution against direct comparison of results from another recent study from a region in England [29], as that study did not take into account prescribing of other antibiotics or the potential differences in the propensity to send in samples from patients. Our results show that areas with low testing rates have artificial high resistance proportions. This finding emphasizes the importance of accounting for differences in testing practices when comparing resistance prevalences between different countries or more granular areas. Ideally sentinel surveillance systems with systematic and standardized testing would be set up to facilitate less biased between-area comparisons and local pre-test resistance probabilities, thereby potentially improving future association studies and clinical practice.

The most obvious limitation of this work is that the associations we found are not necessarily causal. As with any observational study we could not take into account confounding by unmeasured factors, such as antibiotic use in hospitals and other potential selective pressures. The unavailability of hospital prescribing data may have especially affected the analyses focusing on co-amoxiclav and cephalexin as less than half of co-amoxiclav prescribing and approximately half of cephalosporin prescribing occurs in the general practice setting [20]. In addition roughly 40% of fluoroquinolones are prescribed in the hospital setting in England [20]. Such misclassification of exposure/confounders makes the estimated impact of antibiotics that are commonly used in the hospital less reliable. This may partly explain why amoxicillin use – only 13% of penicillins are used in the hospital [20] – is associated with ciprofloxacin resistance, while fluoroquinolone use was not associated with increased amoxicillin prescribing.

Besides the influence of unmeasured confounding, we cannot exclude the possibility that prescribing differs as a consequence of resistance rather than the other way around, i.e. reverse causation. We tried to reduce such reverse causation by looking at antibiotic prescribing happening before the resistance measurement. Nonetheless, reverse causation may reduce the strength of a positive association or even reverse the association. In addition, some of our results may be partly due to other types of model misspecification.

If we were only interested in the influence of one particular antibiotic, a regularization approach that penalizes all coefficients except the antibiotic of interest might provide better estimates [30]. However, because different antibiotics may influence resistance levels in various ways, we decide to penalise all coefficients in the same way, potentially leading to an underestimation of the effect of antibiotics that increase the prevalence of resistance. Therefore, our estimates should be regarded as conservative.

### Conclusion

Amoxicillin prescribing is associated with increased resistance to amoxicillin, ciprofloxacin and trimethoprim. Amoxicillin is the most frequently used antibiotic for respiratory conditions, which are responsible for the largest share in inappropriate antibiotic prescribing in primary care. These findings suggest that there is a potential to reduce selective pressure via (co-)selection with unnecessary use of amoxicillin for viral and self-limiting respiratory tract infection.

Nitrofurantoin prescribing is associated with lower levels of resistance to amoxicillin and trimethoprim, while trimethoprim prescribing is associated with increased levels of amoxicillin, ciprofloxacin and trimethoprim resistance. This suggests that replacing trimethoprim prescribing with nitrofurantoin prescribing where possible for uncomplicated urinary tract infections may also be associated with a reduction in trimethoprim, amoxicillin and ciprofloxacin resistance among *E. coli*. The methodology used in this study, that can cope with correlated antibiotic use, can be used in other settings to further explore the complex relationships between antibiotic use and levels of antibiotic resistance.

## Methods

### Data

All data were collected as part of routine surveillance and were anonymized. Ethics Committee approval was therefore not required. Antibiotic prescribing data were obtained from NHS Digital, who collate for all general practices in England the total number of items that are prescribed and dispensed (http://digital.nhs.uk/). Antibiotic groups were created based on the first five characters of the Anatomical Therapeutic Chemical (ATC) classification system (Fig 1). Antibiotic prescribing was expressed in daily defined doses (DDDs) per 1000 persons per day for each calendar month at the clinical commissioning group (CCG) level. Antibiotics were expressed in DDDs as this at least partly captures the dose and duration of treatment, while this would not be the case when expressing use in terms of items. This is important, because dose and duration has been shown to be an important driver of antibiotic resistance [31-33]. Moreover, using DDDs would facilitate incorporating of hospital prescribing when this data becomes available, as antibiotics used in the hospital are typically expressed in terms of DDDs. CCGs were set up by the Health and Social Care Act 2012 to organize the delivery of NHS services in England. From April 2018, general practices in England belong to one of 209 CCGs.

Reports of *E. coli* isolated from urine samples from general practice patients between April 2014 and January 2016 in England were extracted from PHE’s Second Generation Surveillance System (SGSS) (https://fingertips.phe.org.uk/profile/amr-local-indicators). This national voluntary laboratory surveillance system captures antimicrobial susceptibility data of all microorganisms tested. The database contains laboratory reports supplied electronically by approximately 98% of NHS hospital microbiology laboratories in England. Repeat specimen reports received from the same patient with matching causative agents were excluded if the specimen dates were within 30 days [8]. A 30 day cut-off is often used to distinguish between same and new urinary tract infection episodes. Both samples categorized as intermediate (I) and resistant (R) were treated as being resistant. The following antibiotic susceptibility test results for *E. coli* urine samples were analyzed: amoxicillin, cephalexin, ciprofloxacin, co-amoxiclav and nitrofurantoin. At least 75% of reported *E. coli* urine isolates extracted from SGSS were tested for resistance against these antibiotics; levels of susceptibility testing for other antibiotics were not reported frequently enough for a useful analysis. For each calendar month the number of samples tested for resistance against each antibiotic and the number of samples confirmed as resistant against each antibiotic were measured at the CCG level. Measurements were only included when at least 10 samples and at least 75% of samples were tested for resistance against the antibiotic of interest in the CCG.

### Analyses

Elastic net regularization was used to evaluate the association between the different antibiotic groups and the five resistances of interest [8, 15, 16]. Elastic net regularization combines the advantages of least absolute shrinkage and selection operator (lasso) [17] and ridge regression [18]. Elastic net regularization is especially useful when encountering situations with high collinearity, such as strong correlations in antibiotic usage, and a relatively large number of variables (antibiotic groups) compared to the amount of observations [16, 18, 19]. More conventional regression techniques would likely result in multi-collinearity and sparsity bias issues [18, 19].

We fitted a separate Poisson model with elastic net regularization for each resistance. The number of *E. coli* isolates from urinary samples reported to be resistant each month was included as the dependent variable. The natural logarithm of the number of samples being tested was included as an offset to account for the fact that there is variation in the number of samples tested between CCGs, thereby effectively modelling resistance as a proportion.

Potential explanatory variables were all antibiotics groups (e.g. ATC codes J01AA and J01CA) prescribed in the month before the monthly measured resistance prevalence (expressed in DDDs per 1000 persons per day), month of the year, calendar year and the test rate. The test rate was defined as the number of *E. coli* urinary samples tested for the resistance of interest per 1000 persons-months. The test rate was included because we have previously observed a relatively strong negative relationship between the test rate and the proportion of samples that are resistant [8].Antibiotics and the test rate were standardized by mean-centering and dividing by two standard deviations. To keep the antibiotic groups on the same scale, all antibiotic groups were mean-centered and divided by two standard deviations of penicillins with extended spectrum (ATC code J01CA) instead of using the standard deviations of individual antibiotic groups. To keep all variables, including binary (dummy) variables, on the same scale all variables were divided by two standard deviations [34]. After performing the elastic net regularization variables were back transformed to the original ‘DDD per 1000 persons per day’ scale.

All elastic net analyses were performed using the ‘glmnet’ package in R version 3.4.3 [16]. To reduce the false discovery rate often observed with standard application of regularization methods, we estimated the optimal shrinkage parameter λ using the Akaike information criterion (AIC) [8, 35]. Confidence intervals (CIs) were obtained by taking 1000 clustered bootstrap samples, resampling at the highest level (CCG) with replacement.

### Secondary analysis

In a secondary analysis we varied the lag time between antibiotic prescribing and the resistances of interest. First, instead of using antibiotic prescribing in the month before the resistance measurement as a potential covariate, we evaluated the association between antibiotics used in 1-3 months before the resistance measurements. Further, we assessed the association between antibiotics used in the year (1-12 months) before the resistance measurements.

In addition, we performed a sensitivity analysis, restricting to months with at least 20 measurements to assess the potential influence of potential random error and systematic error due to low sampling rates.

## Funding

SH and JV are supported by the National Institute for Health Research Health Protection Research Unit (NIHR HPRU) in Healthcare Associated Infections and Antimicrobial Resistance at the University of Oxford in partnership with Public Health England (PHE) (HPRU-2012-10041). The views expressed are those of the authors and not necessarily those of the National Health Service, NIHR, Department of Health, or PHE.

**S1 Fig S1. Variation in antibiotic prescribing.**

Data shown for dispensing of tetracyclines, penicillins with extended spectrum, Beta-lactamase-resistant penicillins and macrolide. Each line represents a different clinical commissioning group in England.

**S1 Fig S2. Variation in antibiotic prescribing.**

Data shown for dispensing of trimethoprim, nitrofurantoin, beta-lactamase-sensitive penicillins, and other antibiotics. Each line represents a different clinical commissioning group in England.

**S1 Fig S3. Proportion of urinary samples with *E. coli* isolated resistant to amoxicillin/ampicillin.** The boxplot shows the variation in amoxicillin/ampicillin resistance between Clinical Commissioning Groups over time.

**S1 Fig S4. Proportion of urinary samples with *E. coli* isolated resistant to co-amoxiclav.** The boxplot shows the variation in co-amoxiclav resistance between Clinical Commissioning Groups over time.

**S1 Fig S5. Proportion of urinary samples with *E. coli* isolated resistant to cephalexin.** The boxplot shows the variation in cephalexin resistance between Clinical Commissioning Groups over time.

**S1 Fig S6. Proportion of urinary samples with *E. coli* isolated resistant to ciprofloxacin.** The boxplot shows the variation in ciprofloxacin resistance between Clinical Commissioning Groups over time.

**S1 Fig S7. Proportion of urinary samples with *E. coli* isolated resistant to nitrofurantoin.** The boxplot shows the variation in nitrofurantoin resistance between Clinical Commissioning Groups over time.

**S2 Table S1. Associations between amoxicillin resistance among *E. coli* urinary samples and antibiotic prescribing.**

Antibiotic use is expressed in DDD per 1000 persons per day.

**S2 Table S2. Associations between nitrofurantoin resistance among *E. coli* urinary samples and antibiotic prescribing.**

Antibiotic use is expressed in DDD per 1000 persons per day.

**S2 Table S3. Associations between cephalexin resistance among *E. coli* urinary samples and antibiotic prescribing.**

Antibiotic use is expressed in DDD per 1000 persons per day.

**S2 Table S4. Associations between ciprofloxacin resistance among *E. coli* urinary samples and antibiotic prescribing.**

Antibiotic use is expressed in DDD per 1000 persons per day.

**S2 Table S5. Associations between co-amoxiclav resistance among *E. coli* urinary samples and antibiotic prescribing.**

Antibiotic use is expressed in DDD per 1000 persons per day.

**S2 Table S6. Associations between amoxicillin resistance among *E. coli* urinary samples and antibiotic prescribing using different thresholds of the minimum number of samples tested.**

Antibiotic use is expressed in DDD per 1000 persons per day.

**S2 Table S7. Associations between nitrofurantoin resistance among *E. coli* urinary samples and antibiotic prescribing using different thresholds of the minimum number of samples tested.**

Antibiotic use is expressed in DDD per 1000 persons per day.

**S2 Table S8. Associations between cephalexin resistance among *E. coli* urinary samples and antibiotic prescribing using different thresholds of the minimum number of samples tested.**

Antibiotic use is expressed in DDD per 1000 persons per day.

**S2 Table S9. Associations between ciprofloxacin resistance among *E. coli* urinary samples and antibiotic prescribing using different thresholds of the minimum number of samples tested.**

Antibiotic use is expressed in DDD per 1000 persons per day.

**S2 Table S10. Associations between co-amoxiclav resistance among *E. coli* urinary samples and antibiotic prescribing using different thresholds of the minimum number of samples tested.**

Antibiotic use is expressed in DDD per 1000 persons per day.

