## Supplementary material for "Selection and co-selection of antibiotic resistances among *Escherichia coli* by antibiotic use in primary care: an ecological analysis": S1 Supplementary file

### S1. Supplementary material

#### Variation in antibiotic prescribing

Fig S1 and S2 show the variation in antibiotic prescribing over time in Clinical Commissioning Groups (CCGs) in England. There was generally more variation between CCGs than variation over time within CCGs. However, for some antibiotics there were clear peaks in the amount of dispensed antibiotics. Penicillins with extended spectrum (mainly amoxicillin) are mainly prescribed for respiratory tract infections and prescribing clearly peaked during the winter when the incidence of such infections is higher (peak in December, Fig S1). β-lactamase-resistant penicillins (mainly flucloxacillin) are mainly prescribed for skin conditions and peaked in July in line with the frequently observed summer peak in the incidence of (Gram-positive) skin infections. β-lactamase-sensitive penicillins (mainly penicillin V) are mainly prescribed for sore throat and peaked in March.

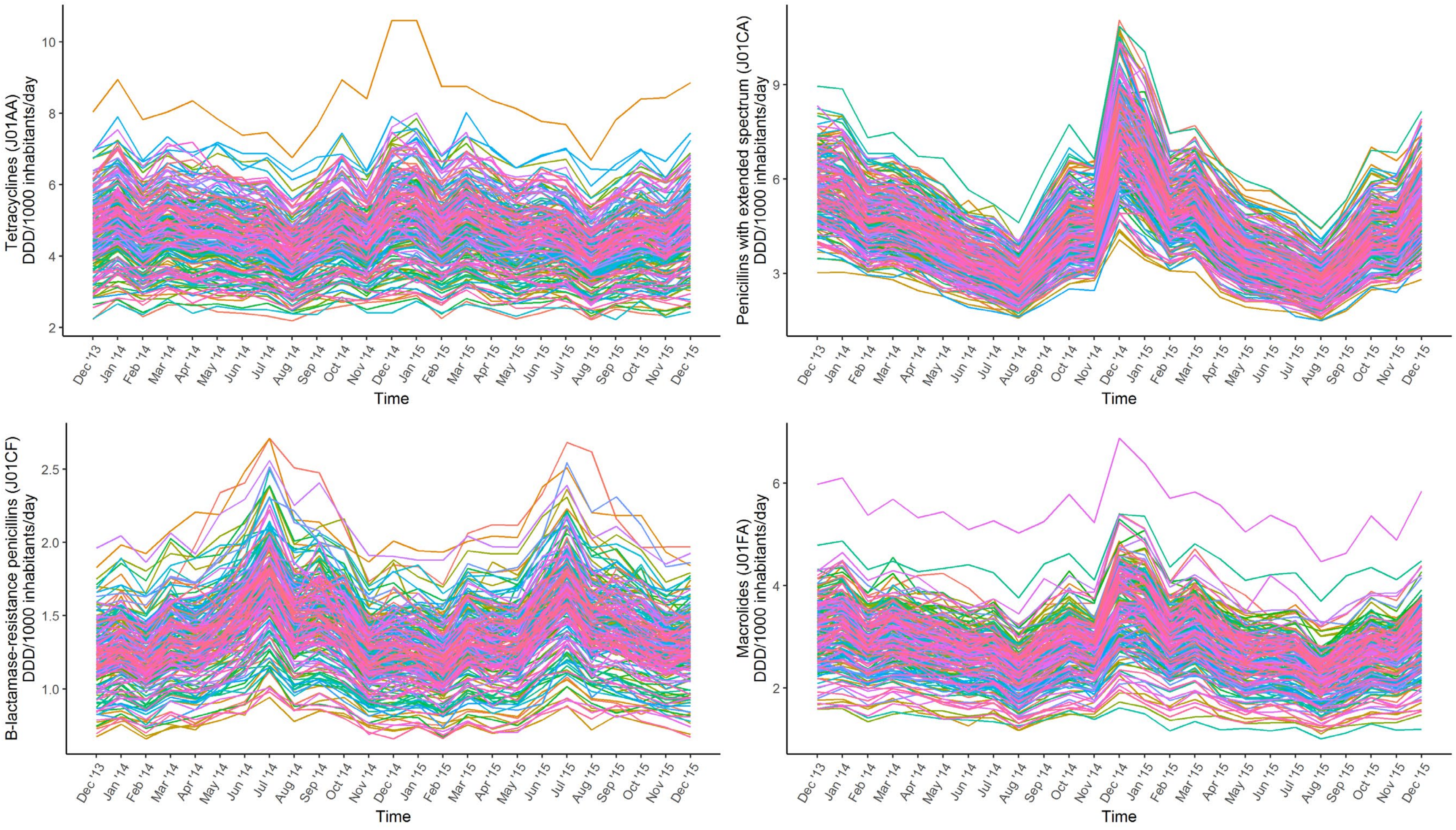


Fig S1. Variation in antibiotic prescribing. Each line represents a different clinical commissioning group in England.

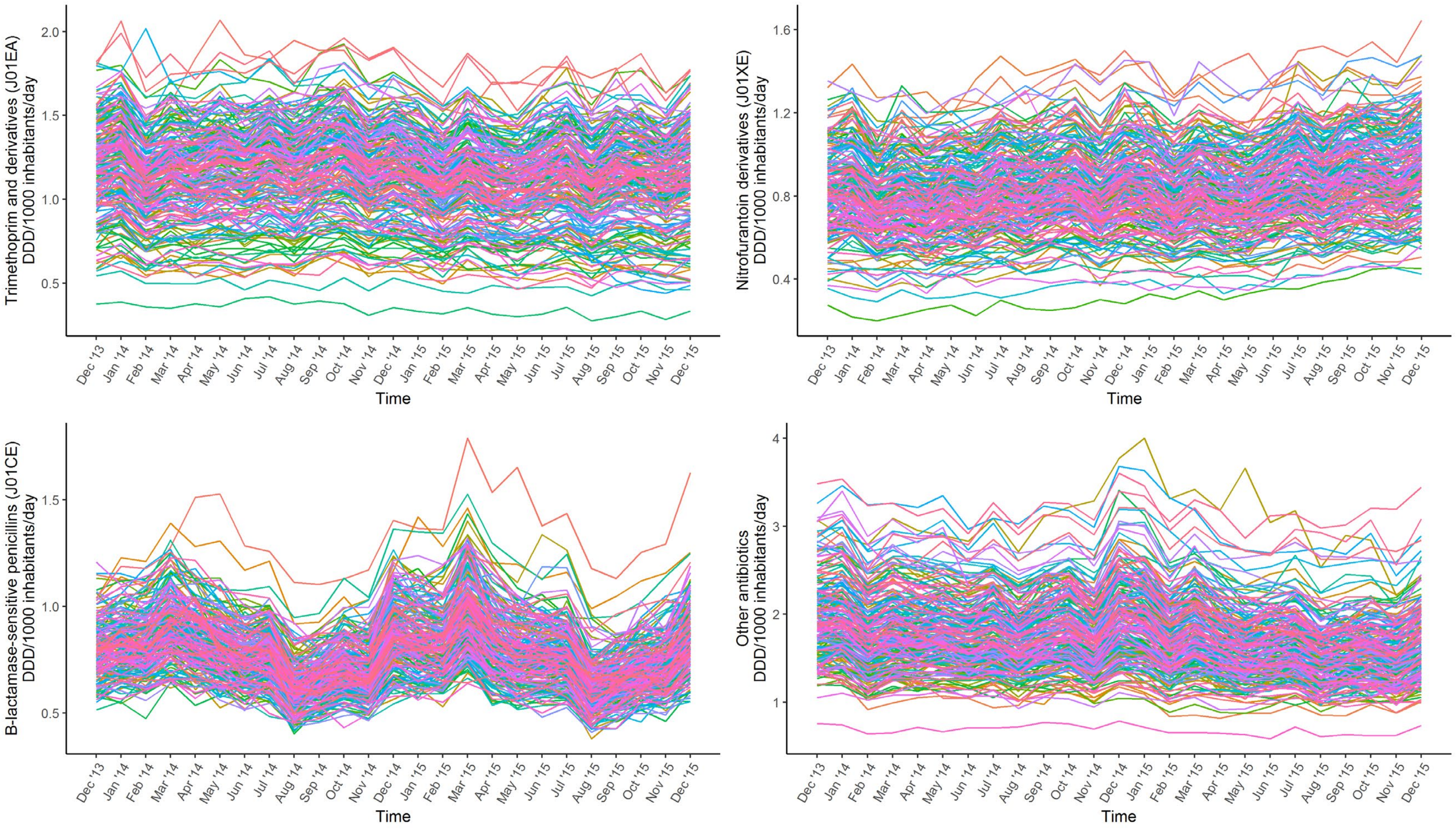


Fig S2. Variation in antibiotic prescribing. Each line represents a different clinical commissioning group in England.

**Variation in antibiotic resistances**

The Figs below show the proportion of *E.coli* urinary samples that is resistant to the antibiotics of interest. The unit of analysis is the Clinical Commissioning Group (CCG). Each month (x-axis) of the boxplot shows the variation across all CCGs, provided that there were sufficient samples tested in that month for the resistance of interest. The Figs show that there is substantial variation between the CCGs (within each month), but that the median (and IQR) proportion of samples being tested as resistant to the antibiotics of interest is more stable over time (between months).

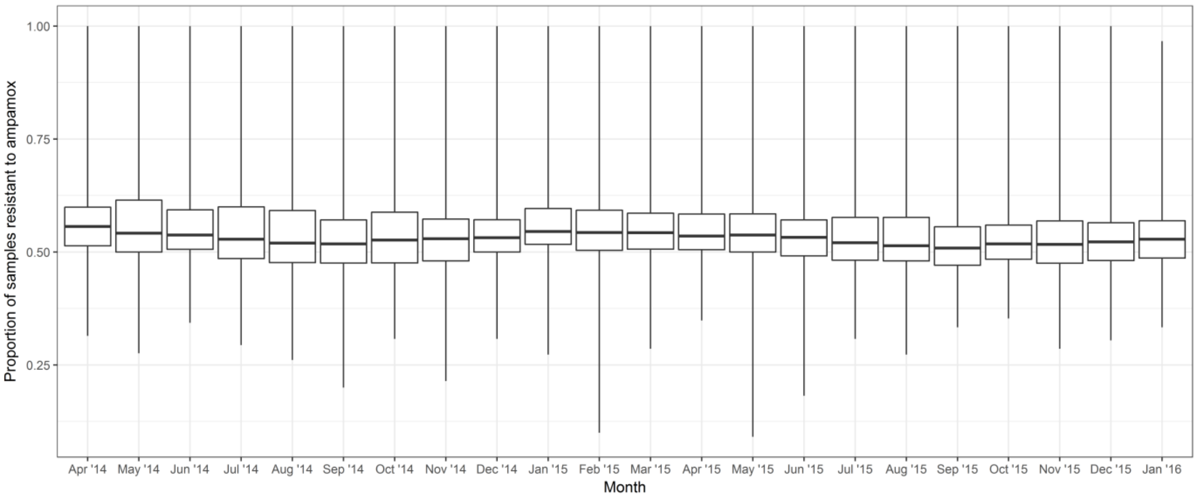


Fig S3. Proportion of urinary samples with *E. coli* isolated resistant to amoxicillin/ampicillin. The boxplot shows the variation in amoxicillin/ampicillin resistance between Clinical Commissioning Groups (geographical regions in England) over time.

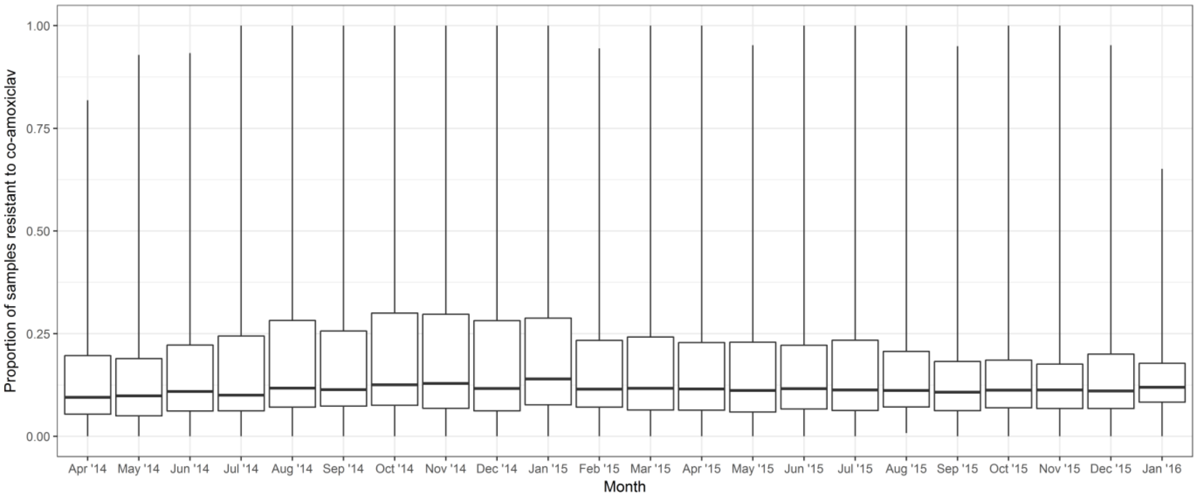


Fig S4. Proportion of urinary samples with *E. coli* isolated resistant to co-amoxiclav. The boxplot shows the variation in co-amoxiclav resistance between Clinical Commissioning Groups (geographical regions in England) over time.

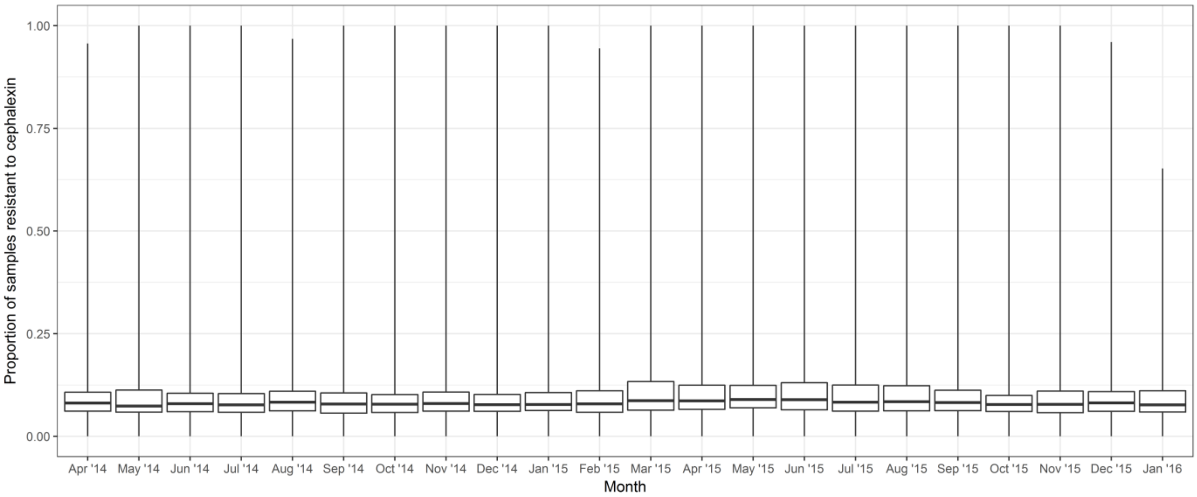


Fig S5. Proportion of urinary samples with *E. coli* isolated resistant to cephalexin. The boxplot shows the variation in cephalexin resistance between Clinical Commissioning Groups (geographical regions in England) over time.

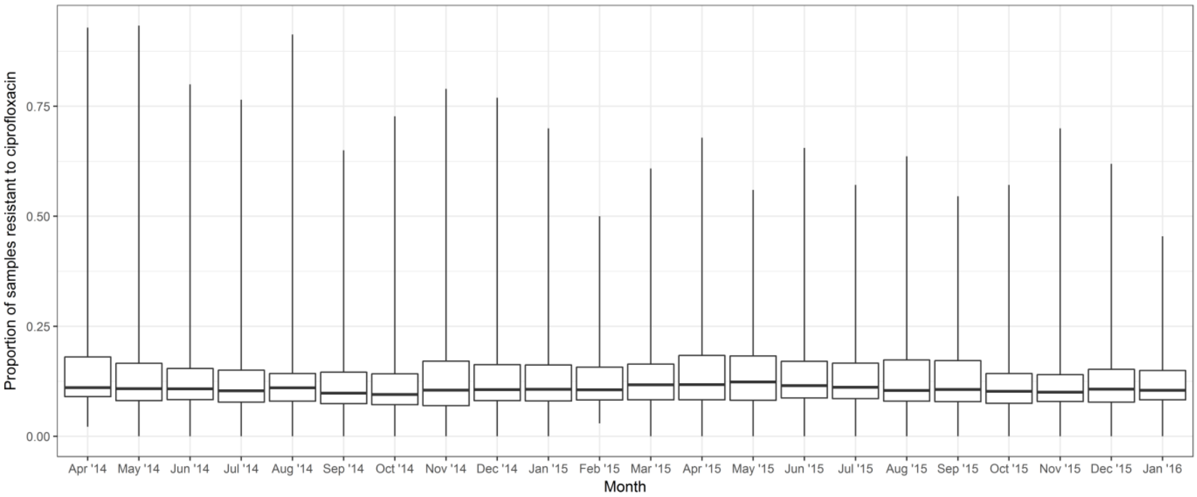


Fig S6. Proportion of urinary samples with *E. coli* isolated resistant to ciprofloxacin. The boxplot shows the variation in ciprofloxacin resistance between Clinical Commissioning Groups (geographical regions in England) over time.

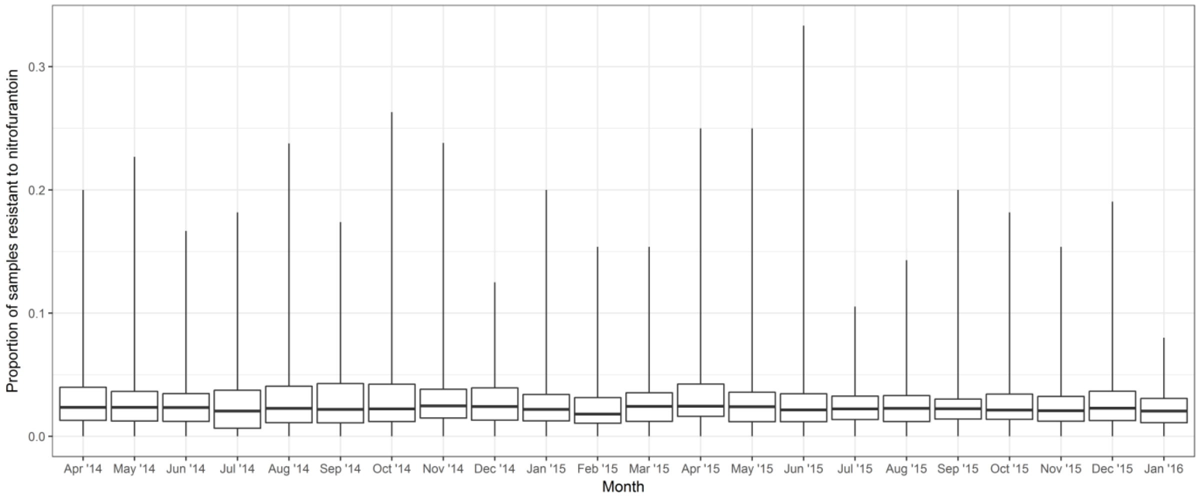


Fig S7. Proportion of urinary samples with *E. coli* isolated resistant to nitrofurantoin. The boxplot shows the variation in nitrofurantoin resistance between Clinical Commissioning Groups (geographical regions in England) over time.
