## Supplementary material for "Selection and co-selection of antibiotic resistances among *Escherichia coli* by antibiotic use in primary care: an ecological analysis": S2 Supplementary file

Tables reporting full model results, including test rate, month and year & Tables reporting results using different thresholds for the minimum number of samples tested in a month.

**Table S1. Associations between amoxicillin resistance among *E. coli* urinary samples and antibiotic prescribing.**

Antibiotic use is expressed in DDD per 1000 persons per day.

|  | Amoxicillin resistance, antibiotic prescribing 1 month before. RR (2.5th–97.5th percentile of bootstrap) |
| --- | --- |
| Tetracyclines (J01AA) | 1.00 (0.98 – 1.01) |
| Penicillins with extended spectrum (J01CA) | 1.03 (1.01 – 1.04) |
| Beta-lactamase-sensitive penicillins (J01CE) | 1.02 (0.97 – 1.12) |
| Beta-lactamase-resistant penicillins (J01CF) | 1.03 (0.98 – 1.12) |
| Combinations of penicillins, including β-lactamase inhibitors (J01CR) | 1.02 (0.95 – 1.08) |
| First-generation cephalosporins (J01DB) | 1.01 (0.91 – 1.07) |
| Second-generation cephalosporins (J01DC) | 1.00 (0.85 – 1.02) |
| Trimethoprim and derivatives (J01EA) | 1.01 (0.98 – 1.08) |
| Macrolides (J01FA) | 0.99 (0.97 – 1.02) |
| Lincosamides (J01FF) | 0.98 (0.71 – 1.00) |
| Fluoroquinolones (J01MA) | 0.93 (0.78 – 0.99) |
| Polymyxins (J01XB) | 1.01 (0.93 – 1.35) |
| Nitrofuran derivatives (J01XE) | 0.92 (0.84 – 0.97) |
| Other antibacterials (J01XX) | 0.98 (0.87 – 1.13) |
| Test rate | 0.97 (0.96 – 0.98) |
| *Year* |  |
| 2014 | Ref. |
| 2015 | 1.00 (1.00 – 1.01) |
| 2016 | 1.01 (1.00 – 1.02) |
| *Month* |  |
| January | Ref. |
| February | 1.00 (0.99 – 1.01) |
| March | 1.01 (1.00 – 1.02) |
| April | 1.01 (1.00 – 1.02) |
| May | 1.01 (1.00 – 1.02) |
| June | 1.01 (1.00 – 1.03) |
| July | 1.01 (0.99 – 1.03) |
| August | 1.00 (0.98 – 1.03) |
| September | 1.01 (0.99 – 1.03) |
| October | 1.01 (0.99 – 1.03) |
| November | 1.00 (0.99 – 1.02) |
| December | 1.01 (1.00 – 1.03) |

**Table S2. Associations between nitrofurantoin resistance among *E. coli* urinary samples and antibiotic prescribing.**

Antibiotic use is expressed in DDD per 1000 persons per day.

|  | Nitrofurantoin resistance, antibiotic prescribing 1 month before. RR (2.5th–97.5th percentile of bootstrap) |
| --- | --- |
| Tetracyclines (J01AA) | 1.01 (0.89 – 1.14) |
| Penicillins with extended spectrum (J01CA) | 1.02 (0.96 – 1.15) |
| Beta-lactamase-sensitive penicillins (J01CE) | 1.50 (0.73 – 2.02) |
| Beta-lactamase-resistant penicillins (J01CF) | 1.01 (0.75 – 1.72) |
| Combinations of penicillins, including β-lactamase inhibitors (J01CR) | - |
| First-generation cephalosporins (J01DB) | 1.76 (0.56 – 3.18) |
| Trimethoprim and derivatives (J01EA) | 1.52 (1.15 – 2.08) |
| Combinations of sulfonamides and trimethoprim, including derivatives (J01EE) | - |
| Macrolides (J01FA) | 0.93 (0.80 – 1.06) |
| Other aminoglycosides (J01GB) | 45.87 (1.00 – 2.12x10^3^) |
| Polymyxins (J01XB) | - |
| Nitrofuran derivatives (J01XE) | 1.52 (1.00 – 2.24) |
| Other antibacterials (J01XX) | 0.84 (0.47 – 1.89) |
| Test rate | 1.01 (0.95 – 1.04) |
| *Year* |  |
| 2014 | Ref. |
| 2015 | 0.95 (0.93 – 0.99) |
| 2016 | 0.91 (0.82 – 1.00) |
| *Month* |  |
| January | Ref. |
| February | 0.98 (0.95 – 1.02) |
| March | 1.06 (1.00 – 1.15) |
| April | 1.00 (0.94 – 1.10) |
| May | 1.02 (0.94 – 1.13) |
| June | 1.02 (0.92 – 1.14) |
| July | 1.00 (0.88 – 1.13) |
| August | 1.02 (0.87 – 1.16) |
| September | 1.05 (0.90 – 1.19) |
| October | 1.02 (0.89 – 1.14) |
| November | 1.00 (0.91 – 1.10) |
| December | 1.02 (0.94 – 1.12) |

**Table S3. Associations between cephalexin resistance among *E. coli* urinary samples and antibiotic prescribing.**

Antibiotic use is expressed in DDD per 1000 persons per day.

|  | Cephalexin resistance, antibiotic prescribing 1 month before. RR (2.5th–97.5th percentile of bootstrap) |
| --- | --- |
| Tetracyclines (J01AA) | 0.97 (0.89 – 1.02) |
| Penicillins with extended spectrum (J01CA) | 0.99 (0.94 – 1.06) |
| Beta-lactamase-sensitive penicillins (J01CE) | 0.98 (0.63 – 1.00) |
| Beta-lactamase-resistant penicillins (J01CF) | 0.98 (0.70 – 1.35) |
| Combinations of penicillins, including β-lactamase inhibitors (J01CR) | 0.98 (0.69 – 1.10) |
| First-generation cephalosporins (J01DB) | 1.01 (0.94 – 1.41) |
| Trimethoprim and derivatives (J01EA) | 1.03 (0.98 – 1.75) |
| Macrolides (J01FA) | 0.95 (0.75 – 1.00) |
| Fluoroquinolones (J01MA) | 1.00 (0.81 – 1.30) |
| Nitrofuran derivatives (J01XE) | 1.01 (0.94 – 1.44) |
| Other antibacterials (J01XX) | - |
| Test rate | 0.84 (0.76 – 0.89) |
| *Year* |  |
| 2014 | Ref. |
| 2015 | 1.02 (1.00 – 1.05) |
| 2016 | 1.01 (0.91 – 1.04) |
| *Month* |  |
| January | Ref. |
| February | 1.01 (0.99 – 1.03) |
| March | 1.02 (0.96 – 1.06) |
| April | 1.02 (0.98 – 1.05) |
| May | 1.00 (0.93 – 1.02) |
| June | 1.00 (0.90 – 1.03) |
| July | 1.00 (0.89 – 1.03) |
| August | 0.99 (0.87 – 1.02) |
| September | 0.98 (0.82 – 1.00) |
| October | 0.98 (0.84 – 0.99) |
| November | 0.98 (0.87 – 0.99) |
| December | 0.98 (0.86 – 1.00) |

**Table S4. Associations between ciprofloxacin resistance among *E. coli* urinary samples and antibiotic prescribing.**

Antibiotic use is expressed in DDD per 1000 persons per day.

|  | Ciprofloxacin resistance, antibiotic prescribing 1 month before. RR (2.5th–97.5th percentile of bootstrap) |
| --- | --- |
| Tetracyclines (J01AA) | 0.92 (0.88 – 0.98) |
| Penicillins with extended spectrum (J01CA) | 1.09 (1.04 – 1.17) |
| Beta-lactamase-resistant penicillins (J01CF) | - |
| Trimethoprim and derivatives (J01EA) | 1.34 (1.10 – 1.59) |
| Macrolides (J01FA) | 0.85 (0.76 – 0.94) |
| Fluoroquinolones (J01MA) | 1.24 (1.00 – 2.81) |
| Nitrofurantoin | - |
| Test rate | 0.83 (0.79 – 0.86) |
| *Year* |  |
| 2014 | Ref. |
| 2015 | 1.01 (0.99 – 1.03) |
| 2016 | 1.00 (0.98 – 1.09) |
| *Month* |  |
| January | Ref. |
| February | 1.00 (0.99 – 1.05) |
| March | 1.02 (1.01 – 1.11) |
| April | 1.04 (1.02 – 1.14) |
| May | 1.03 (1.01 – 1.14) |
| June | 1.03 (1.01 – 1.16) |
| July | 1.03 (1.01 – 1.18) |
| August | 1.01 (0.98 – 1.17) |
| September | 1.01 (0.98 – 1.18) |
| October | 1.00 (0.97 – 1.13) |
| November | 0.99 (0.97 – 1.09) |
| December | 1.00 (0.99 – 1.10) |

**Table S5. Associations between co-amoxiclav resistance among *E. coli* urinary samples and antibiotic prescribing.**

Antibiotic use is expressed in DDD per 1000 persons per day.

|  | Co-amoxiclav resistance, antibiotic prescribing 1 month before. RR (2.5th–97.5th percentile of bootstrap) |
| --- | --- |
| Tetracyclines (J01AA) | 1.03 (0.89 – 1.31) |
| Penicillins with extended spectrum (J01CA) | 0.99 (0.74 – 1.00) |
| Beta-lactamase-sensitive penicillins (J01CE) | - |
| Combinations of penicillins, including β-lactamase inhibitors (J01CR) | - |
| Trimethoprim and derivatives (J01EA) | - |
| Test rate | 0.96 (0.81 – 1.00) |
| *Year* |  |
| 2014 | Ref. |
| 2015 | 0.99 (0.93 – 1.02) |
| 2016 | 1.00 (0.79 – 1.01) |
| *Month* |  |
| January | Ref. |
| February | - |
| March | - |
| April | 1.00 (0.75 – 1.00) |
| May | 1.00 (0.74 – 1.00) |
| June | - |
| July | - |
| August | - |
| September | - |
| October | - |
| November | - |
| December | -- |

**Table S6. Associations between amoxicillin resistance among *E. coli* urinary samples and antibiotic prescribing using different thresholds of the minimum number of samples tested.**

Antibiotic use is expressed in DDD per 1000 persons per day.

| Antibiotic prescribed | Amoxicillin resistance, antibiotic prescribing 1 month before. Restricted to months with ≥20 samples tested. RR (2.5th–97.5th percentile of bootstrap) | Amoxicillin resistance, antibiotic prescribing 1 month before. Main analysis, restricting to months with ≥10 samples tested. RR (2.5th–97.5th percentile of bootstrap) |
| --- | --- | --- |
| Tetracyclines (J01AA) | 1.00 (0.99 – 1.02) | 1.00 (0.98 – 1.01) |
| Penicillins with extended spectrum (J01CA) | **1.02 (1.01 – 1.04)*** | **1.03 (1.01 – 1.04)*** |
| Beta-lactamase-sensitive penicillins (J01CE) | 1.01 (1.00 – 1.15) | 1.02 (0.97 – 1.12) |
| Beta-lactamase-resistant penicillins (J01CF) | 1.02 (0.97 – 1.09) | 1.03 (0.98 – 1.12) |
| Combinations of penicillins, including β-lactamase inhibitors (J01CR) | 1.00 (0.92 – 1.06) | 1.02 (0.95 – 1.08) |
| First-generation cephalosporins (J01DB) | - | 1.01 (0.91 – 1.07) |
| Second-generation cephalosporins (J01DC) | - | 1.00 (0.85 – 1.02) |
| Trimethoprim and derivatives (J01EA) | - | 1.01 (0.98 – 1.08) |
| Macrolides (J01FA) | 1.00 (0.98 – 1.03) | 0.99 (0.97 – 1.02) |
| Lincosamides (J01FF) | - | 0.98 (0.71 – 1.00) |
| Fluoroquinolones (J01MA) | 0.92 (0.74 – 1.00) | **0.93 (0.78 – 0.99)*** |
| Polymyxins (J01XB) | - | 1.01 (0.93 – 1.35) |
| Nitrofuran derivatives (J01XE) | **0.92 (0.84 – 0.97)*** | **0.92 (0.84 – 0.97) *** |
| Other antibacterials (J01XX) | - | 0.98 (0.87 – 1.13) |

*Associations for which 2.5^th^ and 97.5^th^ percentile of the clustered bootstrap are both indicating an increased or decreased risk.

**Table S7. Associations between nitrofurantoin resistance among *E. coli* urinary samples and antibiotic prescribing using different thresholds of the minimum number of samples tested.**

Antibiotic use is expressed in DDD per 1000 persons per day.

| Antibiotic prescribed | Nitrofurantoin resistance, antibiotic prescribing 1 month before. Restricted to months with ≥20 samples tested. RR (2.5th–97.5th percentile of bootstrap) | Nitrofurantoin resistance, antibiotic prescribing 1 month before. Main analysis, restricting to months with ≥10 samples tested. RR (2.5th–97.5th percentile of bootstrap) |
| --- | --- | --- |
| Tetracyclines (J01AA) | 1.01 (0.89 – 1.14) | 1.01 (0.89 – 1.14) |
| Penicillins with extended spectrum (J01CA) | 1.02 (0.96 – 1.16) | 1.02 (0.96 – 1.15) |
| Beta-lactamase-sensitive penicillins (J01CE) | 1.52 (0.78 – 2.14) | 1.50 (0.73 – 2.02) |
| Beta-lactamase-resistant penicillins (J01CF) | - | 1.01 (0.75 – 1.72) |
| First-generation cephalosporins (J01DB) | 1.77 (0.57 – 2.97) | 1.76 (0.56 – 3.18) |
| Trimethoprim and derivatives (J01EA) | **1.51 (1.13 – 2.05)*** | **1.52 (1.15 – 2.08)*** |
| Macrolides (J01FA) | 0.94 (0.79 – 1.06) | 0.93 (0.80 – 1.06) |
| Other aminoglycosides (J01GB) | 24.47 (1.00 – 2.08x10^3^ | 45.87 (1.00 – 2.12x10^3^) |
| Nitrofuran derivatives (J01XE) | **1.52 (1.05 – 2.25)*** | **1.52 (1.00 – 2.24)*** |
| Other antibacterials (J01XX) | 0.85 (0.49 – 2.36) | 0.84 (0.47 – 1.89) |

*Associations for which 2.5^th^ and 97.5^th^ percentile of the clustered bootstrap are both indicating an increased or decreased risk.

**Table S8. Associations between cephalexin resistance among *E. coli* urinary samples and antibiotic prescribing using different thresholds of the minimum number of samples tested.**

Antibiotic use is expressed in DDD per 1000 persons per day.

| Antibiotic prescribed | Cephalexin resistance, antibiotic prescribing 1 month before. Restricted to months with ≥20 samples tested. RR (2.5th–97.5th percentile of bootstrap) | Cephalexin resistance, antibiotic prescribing 1 month before. Main analysis, restricting to months with ≥10 samples tested. RR (2.5th–97.5th percentile of bootstrap) |
| --- | --- | --- |
| Tetracyclines (J01AA) | 0.98 (0.91 – 1.02) | 0.97 (0.89 – 1.02) |
| Penicillins with extended spectrum (J01CA) | 0.98 (0.93 – 1.06) | 0.99 (0.94 – 1.06) |
| Beta-lactamase-sensitive penicillins (J01CE) | 0.99 (0.63 – 1.06) | 0.98 (0.63 – 1.00) |
| Beta-lactamase-resistant penicillins (J01CF) | 0.98 (0.66 – 1.18) | 0.98 (0.70 – 1.35) |
| Combinations of penicillins, including β-lactamase inhibitors (J01CR) | 0.99 (0.69 – 1.11) | 0.98 (0.69 – 1.10) |
| First-generation cephalosporins (J01DB) | 1.00 (0.89 – 1.40) | 1.01 (0.94 – 1.41) |
| Trimethoprim and derivatives (J01EA) | 1.00 (0.95 – 1.58) | 1.03 (0.98 – 1.75) |
| Macrolides (J01FA) | 0.96 (0.74 – 1.00) | 0.95 (0.75 – 1.00) |
| Fluoroquinolones (J01MA) | - | 1.00 (0.81 – 1.30) |
| Nitrofuran derivatives (J01XE) | 1.00 (0.97 – 1.49) | 1.01 (0.94 – 1.44) |

*Associations for which 2.5^th^ and 97.5^th^ percentile of the clustered bootstrap are both indicating an increased or decreased risk.

**Table S9. Associations between ciprofloxacin resistance among *E. coli* urinary samples and antibiotic prescribing using different thresholds of the minimum number of samples tested.**

Antibiotic use is expressed in DDD per 1000 persons per day.

| Antibiotic prescribed | Ciprofloxacin resistance, antibiotic prescribing 1 month before. Restricted to months with ≥20 samples tested. RR (2.5th–97.5th percentile of bootstrap) | Ciprofloxacin resistance, antibiotic prescribing 1 month before. Main analysis, restricting to months with ≥10 samples tested. RR (2.5th–97.5th percentile of bootstrap) |
| --- | --- | --- |
| Tetracyclines (J01AA) | **0.92 (0.88 – 0.98)** | **0.92 (0.88 – 0.98)*** |
| Penicillins with extended spectrum (J01CA) | **1.09 (1.04 – 1.17)** | **1.09 (1.04 – 1.17)*** |
| Trimethoprim and derivatives (J01EA) | **1.32 (1.08 – 1.58)** | **1.34 (1.10 – 1.59)*** |
| Macrolides (J01FA) | **0.84 (0.77 – 0.95)** | **0.85 (0.76 – 0.94)*** |
| Fluoroquinolones (J01MA) | 1.35 (1.00 – 2.87) | 1.24 (1.00 – 2.81) |
| Other antibacterials (J01XX) | 1.11 (0.83 – 4.42) | - |

*Associations for which 2.5^th^ and 97.5^th^ percentile of the clustered bootstrap are both indicating an increased or decreased risk.

**Table S10. Associations between co-amoxiclav resistance among *E. coli* urinary samples and antibiotic prescribing using different thresholds of the minimum number of samples tested.**

Antibiotic use is expressed in DDD per 1000 persons per day.

| Antibiotic prescribed | Co-amoxiclav resistance, antibiotic prescribing 1 month before. Restricted to months with ≥20 samples tested. RR (2.5th–97.5th percentile of bootstrap) | Co-amoxiclav resistance, antibiotic prescribing Main analysis, restricting to months with ≥10 samples tested. RR (2.5th–97.5th percentile of bootstrap) |
| --- | --- | --- |
| Tetracyclines (J01AA) | - | 1.03 (0.89 – 1.31) |
| Penicillins with extended spectrum (J01CA) | - | 0.99 (0.74 – 1.00) |

*Associations for which 2.5^th^ and 97.5^th^ percentile of the clustered bootstrap are both indicating an increased or decreased risk.
